# Potential risk for hearing from prolonged exposure to sound at conversation levels

**DOI:** 10.64898/2026.02.26.708062

**Authors:** Wenyue Xue, Nolan Sun, Emily Wood, Jason Xie, Xiuping Liu, Jun Yan

## Abstract

Prolonged exposure to loud and moderate noise impairs hearing; the lower the noise level, the lower risk of hearing loss is. To date, little is known about how low the noise level can be safe to hearing. This study investigated the risk of exposure to tone at typical conversational levels by measuring auditory brainstem response (ABR). We show that exposing C57 mice to continuous pure tone at 65 dB SPL for 1 hour (TE_65_) leads to an increase in ABR threshold that is specific to the exposure frequency. Tone exposure also increased the latencies and decreased the amplitude in Waves I and II but not in Waves III and V. Significantly, the changes in amplitude and latency were highly correlated in Wave I and such correlation gradually degraded from Wave I through to Wave V. Our findings suggest that exposure to low level sound can impair hearing and alter the auditory information process in the brain if it is persistent and presented over a sufficient period of time.

**Significant Statement:** Our findings established the risk of hearing impairment following the exposure to continuous tone at normal or conversational voice levels. This finding challenges current public health guidelines for hearing protection. Although further clarification is required, our studies prompt that the regular use of ABR testing is a potential protocol for diagnosing hearing impairment in patients experiencing hidden hearing loss (HHL).

## Introduction

Sound, as a media of daily communication, can be hazardous to hearing depending on its intensity and exposure time. A specific type of acquired sensorineural hearing loss known as HHL has attracted increasing attention (Ruggles et al., 2011; Bharadwaj and Shinn-Cunningham, 2021; Liu et al., 2024). HHL refers to hearing difficulty experienced by individuals who show normal hearing sensitivity on an audiogram test and have no history of exposure to loud sound (>105 dB SPL) or drug use (Kohrman et al., 2020; Monaghan et al., 2020). Prolonged exposure to low-level sound is believed to be one of the causes of HHL (Shi et al., 2016; Valderrama et al., 2018).

Exposure to loud sound over sufficient time can cause temporal or permanent hearing loss (decreased sensitivity or increased hearing threshold). Sounds at or below 80 dBA for 8 hours per day (Leq,8 h = 80 dBA) are generally considered safe. However, prolonged exposure to moderate sound (Leq,8 h= 80 dB SPL to Leq,2 h = 105 dB SPL) is postulated to cause HHL and has been studied in animal models for some time (Noreña et al., 2006; Shi et al., 2015; Shi et al., 2016; Liu et al., 2019). Multiple studies reveal that long exposure to moderate sound leads to significant changes in the peripheral and central auditory system with a recoverable hearing threshold (Robertson, 1983; Shi et al., 2013; Singer et al., 2013). For example, exposing mice to noise Leq,2 h = 95-98 dB SPL results in progressive synaptopathy, i.e., the loss of presynaptic ribbons and swelling of terminals of the auditory nerves in the inner ear (Zhao et al., 2021). Functionally, exposing rats to Leq,2 h = 97 dB SPL increases hearing threshold by up to16 dB (Morgan et al., 2019). Significant alterations in neural information processing are also observed in the auditory cortex and midbrain (Noreña et al., 2008; Pienkowski et al., 2011).

The question raised here centers on the impact of exposure to repetitive or prolonged sound at the acoustic level in the living environmental, such as the noise experienced inside a moving car (typically 70-90 dBA) or by the use of personal listening devices (recommended level of <70 dBA) (Flamme et al., 2012; Fink, 2024). To date, only scattered reports have investigated the impact of sound levels below 80 dB SPL. Zhou and Merzenich reported that exposure mice to noise bursts (Leq,24 h = 65 dB SPL) for 2 months leads to a decreased performance score in temporal rate discrimination tasks and with no threshold increase in ABR (Zhou and Merzenich, 2012). Lau and colleagues, by using functional magnetic resonance imaging, demonstrated a level-dependent decrease in the blood oxygenation in the rat medial geniculate body and auditory cortex after a 2-month broadband noise exposure (Leq,24 h = 65 dB SPL). On the other hand, Liu and colleagues recently reported a significant elevation in the threshold of cochlear compound action potential of rats following the exposure to a banded continuous noise (18-24 kHz) at 65 dB SPL for 6 weeks without a significant change in the function of the inferior colliculus (Liu et al., 2021). These studies prompt our study that the exposure to ‘safe’ sound can potentially induce HHL.

To clarify the impact of ‘safe’ sound on hearing, we examined the changes in mouse ABR following the exposure to a 1-hour persistent tone at a level of 65 dB SPL (TE_65_). Our work showed a significant increase in ABR thresholds, a decrease in the amplitude of Wave I (as represented in the activity of the auditory nerve and cochlear nucleus), and finally, the down-regulated correlation of changes in latency vs. amplitude from Wave I through to Wave V.

## Methods

### Animals and Anesthetics

The mice (C57BL/6, 5-7 weeks old, 16.8-21.5g) used in this study were obtained from the Charles River Laboratories in Canada. They were cared for at the Animal Resource Centre, the University of Calgary (UofC) and housed in cages where they were exposed to a 12-hour light/dark cycle and given free access to water and food. The experimental protocol followed was approved by the Animal Care Committee (UofC, Protocol No.: AC22-0115). Mice with a hearing threshold higher than 40 dB SPL were excluded. A total of 30 female mice were used. The mice were anesthetized with a mixture of 85 mg/kg ketamine (Vétoquinol N.-A. Inc, Canada) and 15 mg/kg xylazine (Vétoquinol N.-A. Inc, Canada) through intraperitoneal injection. The anesthetic level was maintained throughout the physiological experiment including the period of tone exposure. The anesthetic level was monitored every 20 minutes by paw pinching. Additional dosages of ketamine (17 mg/kg) and xylazine (3 mg/kg) was administered when mouse showed any withdrawal responses. A head holder was used to fix the mouse’s head by rigidly clamping the palate and the nasal/frontal bones, and the head was adjusted to align the bregma and lambda on one horizontal plane. The mouse’s body temperature was maintained at ∼37°C through a feedback-controlled heating pad. All physiological experimentation and tone exposures were performed in an echo-attenuation soundproof chamber.

### Acoustic Stimulation and Tone Exposure (TE_65_)

Tone bursts (10 ms long with 0.5-ms rising/decay time) were used as acoustic stimuli for the sampling ABR and a 1-hour continuous tone at 65 dB (TE_65_) was used for sound exposure. These acoustic signals were digitally generated and converted into analog signals using an Enhanced Real-time Processor (RP2, Tucker–Davis Tech., Gainesville, FL, USA). Tone was delivered via a digital attenuator (PA5, Tucker–Davis Tech., Gainesville, FL, USA) to a loudspeaker placed 20 cm away, 45 degrees lateral to the mouse’s right ear. The output of the loudspeaker, calibrated for the location corresponding to the right ear of the mouse, is expressed as the decibel sound pressure level (dB SPL) within 1 decibel of accuracy (reference 20 μPa).

### Tone exposure (TE_65_)

Under anesthesia, mice were exposed to a continuous pure tone for 1 hour. The tone was delivered to the right ear of the mouse through the loudspeaker. The frequency of pure tone was set at the characteristic frequency that was determined by the ABR receptive field (see below), and its level was set at 65 dB SPL (a typical conversation level). To ensure unilateral exposure, warm saline drops were applied to block the contralateral ear canal prior to the tone delivery and the ear was cleaned after the exposure.

### ABR Recording

To record the ABR, two stainless steel electrodes were placed subcutaneously, one at the vertex (∼1 mm posterior to the lambda point) and the other at the mastoid (just below the pinna of the right ear). Bioelectrical signals were amplified 5,000 times, filtered with a bandpass of 0.25-2.5 kHz and then fed to an Enhanced Real-time Processor (RP2, Tucker–Davis Tech., Gainesville, FL, USA) for data acquisition (BrainWare software, Tucker–Davis Tech., Gainesville, FL, USA). Tone-evoked ABR potentials were sampled in a time widow of 10 ms starting from the tone onset and ABR data were obtained by averaging 500 traces in response to identical tone.

The ABR receptive field was achieved by recording ABR in response to a set of tones with various frequencies and levels, and saved digitally as a DAM file. The frequencies ranged from 2.5 kHz to 40 kHz with an increment of 0.5 octave and the levels ranged from 15 to 50 dB SPL with an increment of 5 dB plus 50 to 90 dB SPL with an increment of 10 dB. The frequencies and levels of a tone set were presented randomly during the recording with an interval of 50 ms. The ABR receptive fields were recorded twice in each mouse, ie., before and immediately after the TE_65_ (n = 20). To observe the time course of recovery after the TE_65_, the ABR receptive fields were recorded before TE65 and 1, 3, 4, 5, and 6 hours after the TE_65_ using 10 additional mice.

### Data processing

The ABR data stored in the DAM file were read out and displayed by using SoundCode, a custom-made data processing software that allowed us to measure an individual ABR waveform and to characterize the ABR receptive field including characteristic frequency and minimum threshold. Recording traces contaminated by artifacts were all excluded from the trace averaging for delineating the ABR wave. Wave peak latencies were used to identify different waves. Four waves were typically identified in our recordings. Wave III presented as a merged wave III/IV. Waves I-II were prominent and used to determine the characteristic frequency and response threshold based on the ABR receptive field delineated by the series of ABRs in response to the tone set (with various frequencies and levels) as described above.

The minimum threshold of the ABR was defined as the lowest response threshold across all frequencies. The characteristic frequency of the ABR was defined as the frequency at which the minimum threshold was achieved. Wave latency was the range from tone onset to wave peak time. Wave amplitude was the peak voltage subtracted by the voltage from the subsequent valley. Individual ABR waves were identified according to wave peak latencies in responses to the tone of CF and 60 dB SPL; specifically, they were 1.5-2.4 ms for Wave I, 2.5-3.8 ms for Wave II, 3.9-5.5 ms for Wave III/VI, and 5.5-7.5ms for Wave V (Scimemi et al., 2014).

### Statistical analysis

All data were expressed as mean ± standard deviation (SD). A paired t-test (two-tailed) was applied to compare the differences in thresholds and wave amplitudes between pre and post TE_65_ (control vs. TE_65_). Pearson’s correlation coefficient was calculated to assess the linear relationship of the changes between wave amplitude and latency. A t-test (two-tailed) was used to test the significance of the *R*-value of a correlation. A p-value of less than 0.05 was considered statistically significant.

## Results

Tone-induce ABR waveforms pre and post TE_65_ were similar in all mice and typically showed 4 distinct waves within a time window of 10 ms when the tone level was at 60 dB SPL or 70 dB SPL. Following a decrease in the tone level, the number of visible waves also decreased. Wave I/II showed the greatest amplitude response and remained identifiable at the threshold level. The wave amplitudes decreased along with increased latencies following a decrease in the tone level (Fig. 1). The characteristic frequency of the ABR was either 10 kHz (n = 17) or 14.142 kHz (n = 13). The minimum threshold was 22.67 ± 5.37 dB SPL (n = 30) and fell within a range of 15 to 30 dB SPL.

**Figure 1.**
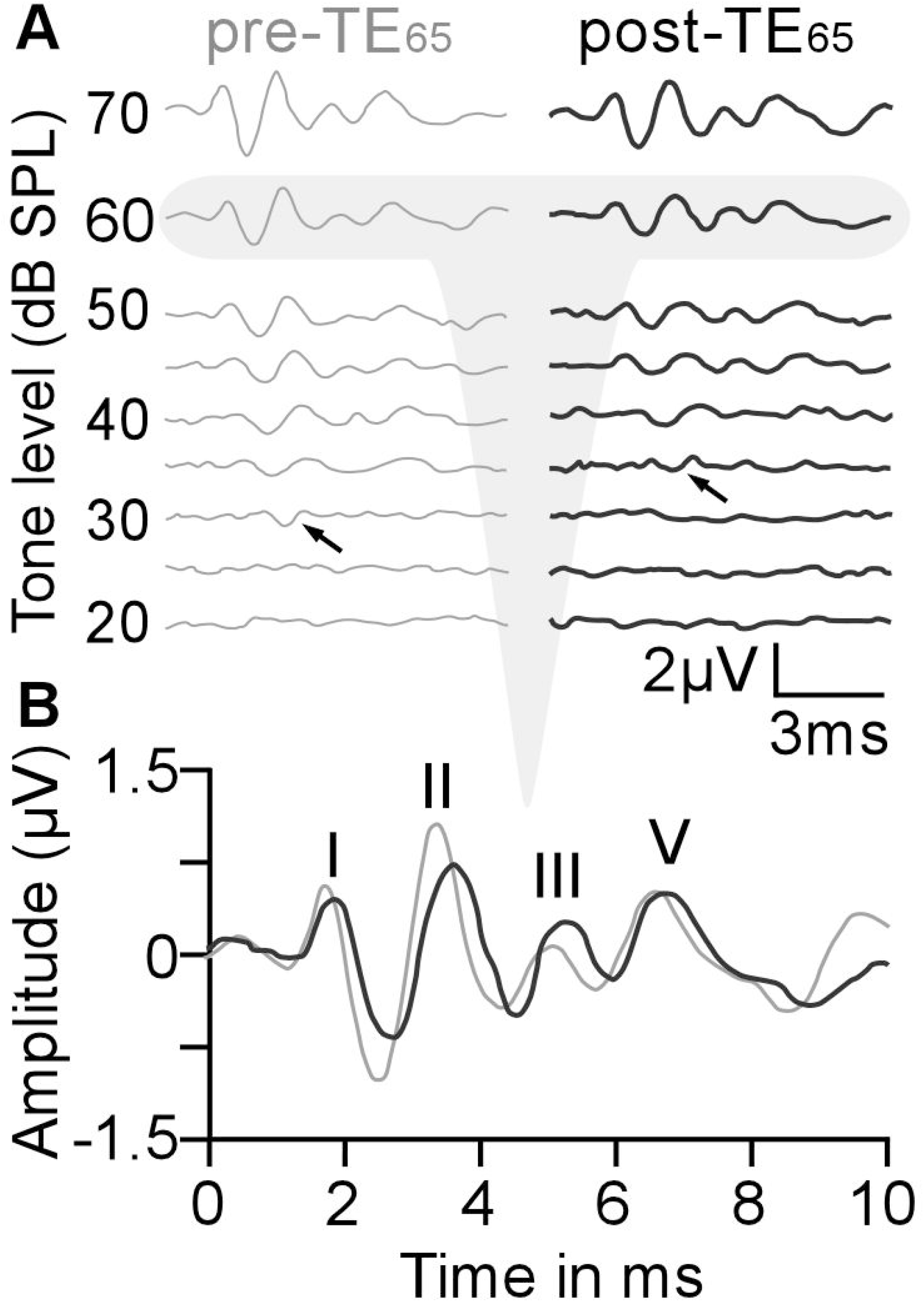
A typical example of an ABR waveform in response to tones with various frequency and levels (**A**) and ABR waves in response to a tone at 60 dB SPL (**B**). Threshold increases following TE_65_ are indicated by black arrows in pre-TE_65_ (grey) and post-TE_65_ (black). Waves I, II, III, and V were identifiable in this ABR waveform.

### ABR threshold increase and its frequency-specificity

The TE_65_ increased the threshold in response to the exposure frequency in 18 out of 30 mice and the largest threshold increase was 15 dB. On average, the threshold increased by 6.39 ± 2.92 dB, from 21.39 ± 5.09 dB SPL to 27.78 ± 6.24 dB SPL (p < 0.001, n = 18). It is important to note here that TE_65_ significantly decreased the amplitude of the ABR wave at the level of the minimum threshold for mice with an unchanged threshold (0.28 ± 0.09 µV vs. 0.22 ± 0.10 µV, p = 0.008, n = 12). At the minimum threshold, one wave was typically identifiable, and this wave mostly represented the Wave I in our recording. To illustrate the TE_65_-impacted areas in relation to the exposure frequency, the ABR frequency tuning curves from all mice were aligned to the exposure frequency as shown in Fig. 2A. TE_65_ appeared to increase the threshold not only at the exposure frequency but also at the frequencies higher than the exposure frequencies in a range of about 1 octave (Fig. 2B). The most significant threshold increase was observed at 0.5 octave above the exposure frequency (26.95 ± 6.67 dB SPL vs. 31.94 ± 6.22 dB SPL, p < 0.0001, n = 18). On the other hand, TE_65_ had little impact on the threshold in response to the frequencies below the exposure frequency (Fig. 2, A and B).

**Figure 2.**
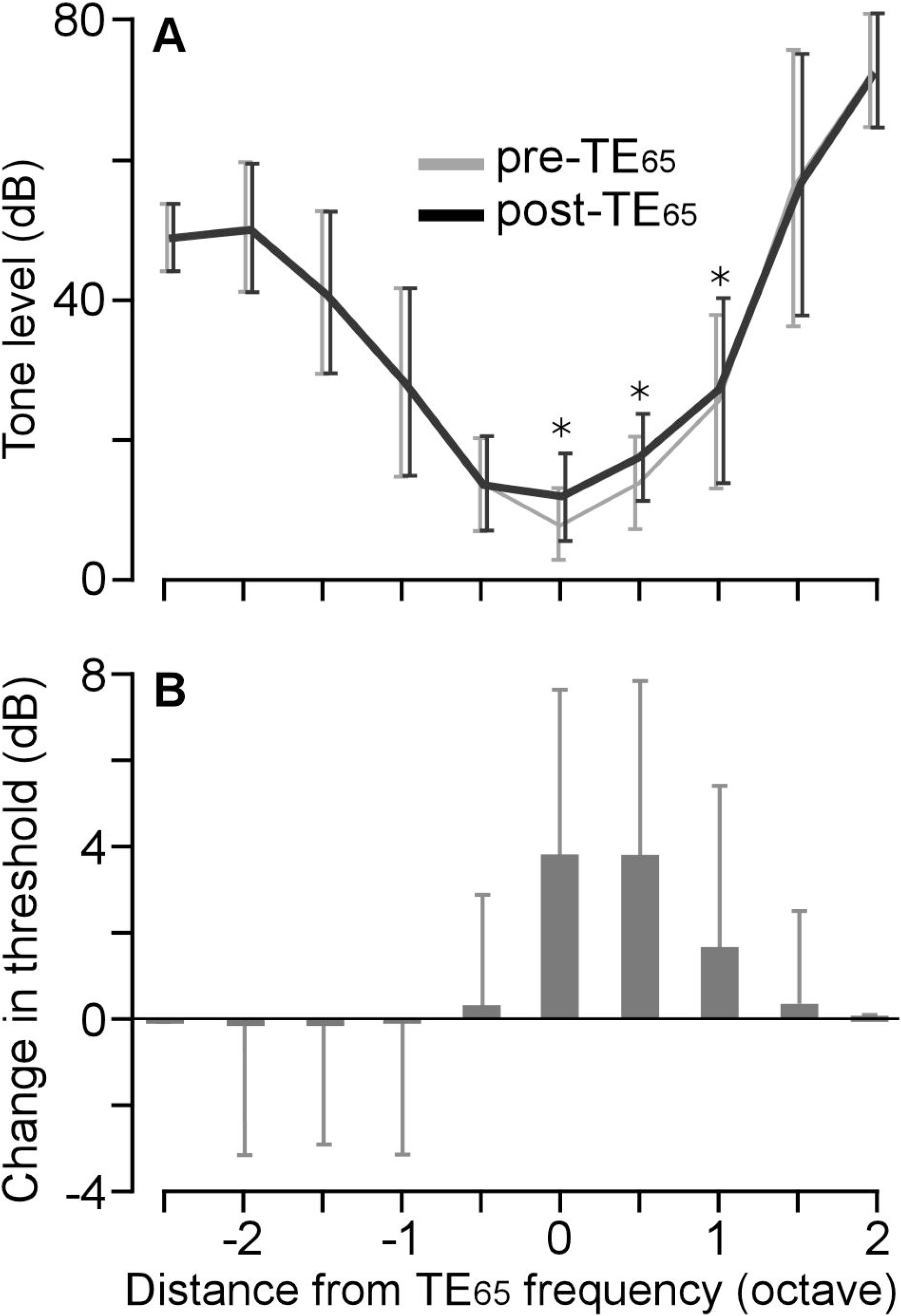
Frequency-dependent changes in ABR thresholds. **A.** A comparison of the averaged ABR thresholds in response to the tones of different frequencies before and after TE_65_. Because the data are aligned to the TE_65_ frequency during averaging, that frequency is shown as 0 octave, and all other frequencies are displayed as their octave distance from TE_65_. **B**. The differences in the thresholds across all frequencies before and after TE_65_. The thresholds in a range from TE_65_ frequency to 1 octave above are remarkably increased by TE_65_.

### Decrease in amplitude and increase in latency of ABR waves

In general, the amplitudes of Waves I, II, III and V changed as the function of tone level, the higher the tone level, the greater the impact on the amplitude. TE_65_ also largely altered the amplitudes of ABR waves, and the Wave I/II amplitudes were always decreased as shown in Figure 1B. In all 30 samples including those with and without threshold increase, the level-amplitude function was remarkably sharper in Waves I and II than in Waves III and V (Fig. 3A). TE_65_ did not alter the overall pattern of level-amplitude function (Fig. 3B). We next analyzed the TE_65_- induced changes in the amplitudes of all waves as the function of tone levels. The changes in Wave amplitudes were expressed as a percentage [100 × (pre-TE_65_ – post-TE_65_)/pre-TE_65_]. TE_65_ demonstrably induced changes in the amplitudes of all four waves as shown by the “U” shape in the range of 20-90 dB SPL (Fig. 3C). The changes of Waves I-II were mostly below 0, indicating an amplitude decrease at all tone levels. The decrease was greatest in the range of 45-60 dB SPL, which represents more than a 20% decrease and the maximum decrease measured 28.5% at 60 dB SPL. With regards to Wave III, the amplitude decreases greater than 20% were observed at 35 dB SPL and 45 dB SPL while the amplitude increases were observed at the levels of 20-, 25- and 90-dB SPL. In contrast, the TE_65_- induced changes in Wave V amplitude fell in the region above 0. This suggests that TE_65_ led to an overall increase in the amplitudes of Wave V. This was an interesting phenomenon considering the general amplitude decreases in Waves I, II and III.

**Figure 3.**
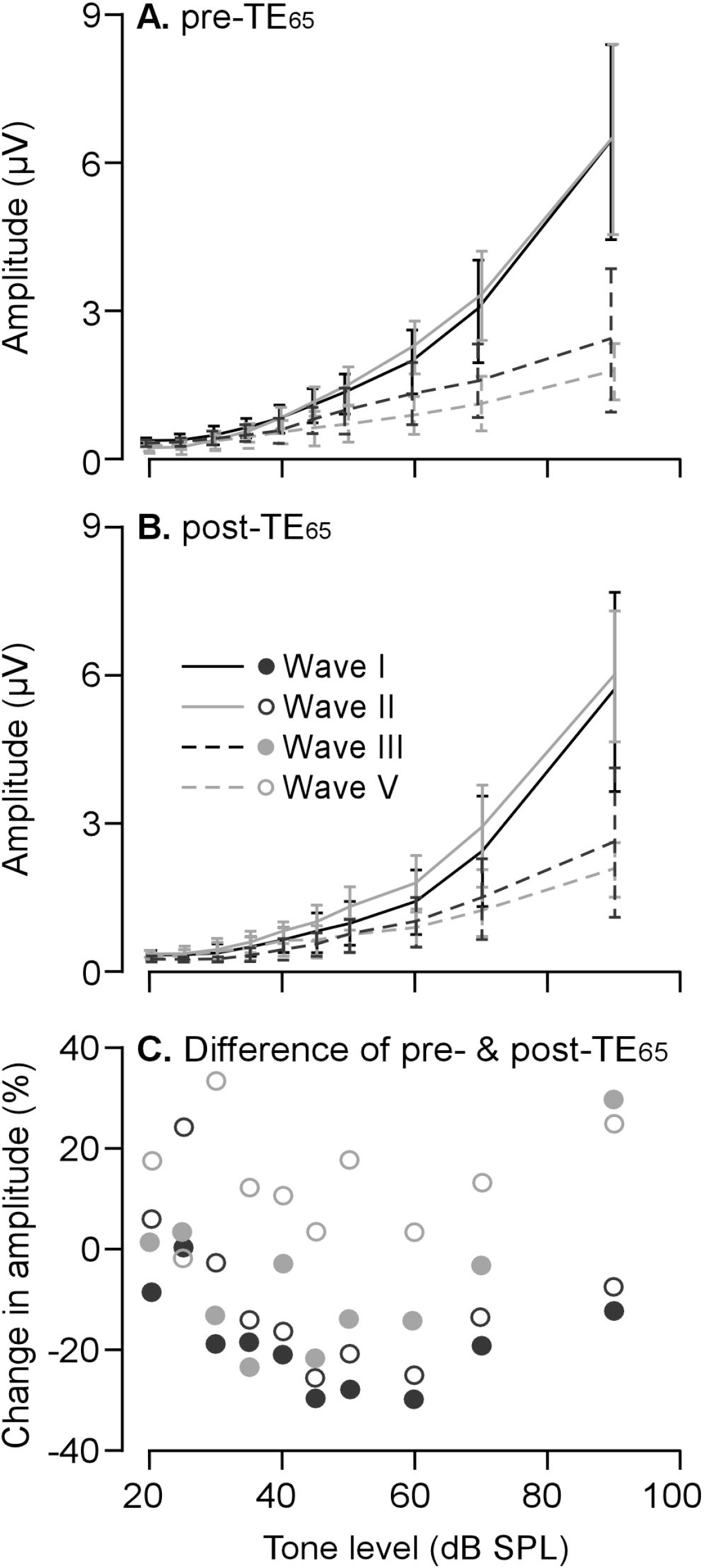
TE_65_-induced changes in the amplitudes across tone levels. The amplitudes of ABR waves before (**A**) and after (**B**) TE_65_ are plotted as the function of tone levels. They show that the patterns of amplitude-level functions were similar before and after TE_65_. The remarkable decrease in the amplitudes of Waves I-III occurred when the tone levels fell within a range of 45 to 60 dB SPL (**C**).

As the greatest decrease in the amplitude of Waves I and II at a tone level of 60 dB SPL, which was based on our analysis of level-amplitude function, we next compared the latency and amplitude of different waves at 60 dB SPL before and after TE_65_. As shown in Figure 4A, TE_65_ significantly decreased the amplitude from 2.14 ± 0.69 µV to 1.52 ± 0.67 µV in Wave I (p < 0.001), from 2.24 ± 0.54 µV to 1.68 ± 0.54 µV in Wave II (p < 0.001), and from 1.23 ± 0.57 µV to 0.98 ± 0.45 µV in Wave III (p < 0.001). The amplitude of Wave V was not significantly changed by the TE_65_. In contrast to wave amplitude changes, TE_65_ significantly increased the latencies of all waves (Fig. 4B). The latency was increased by 0.09 ± 0.27 ms in Wave I (p < 0.001), by 0.16 ± 0.47 ms in Wave II (p < 0.001), by 0.14 ± 0.47 ms in Wave III (p < 0.05), and by 0.26 ± 0.80 ms in Wave V (p < 0.05).

**Figure 4.**
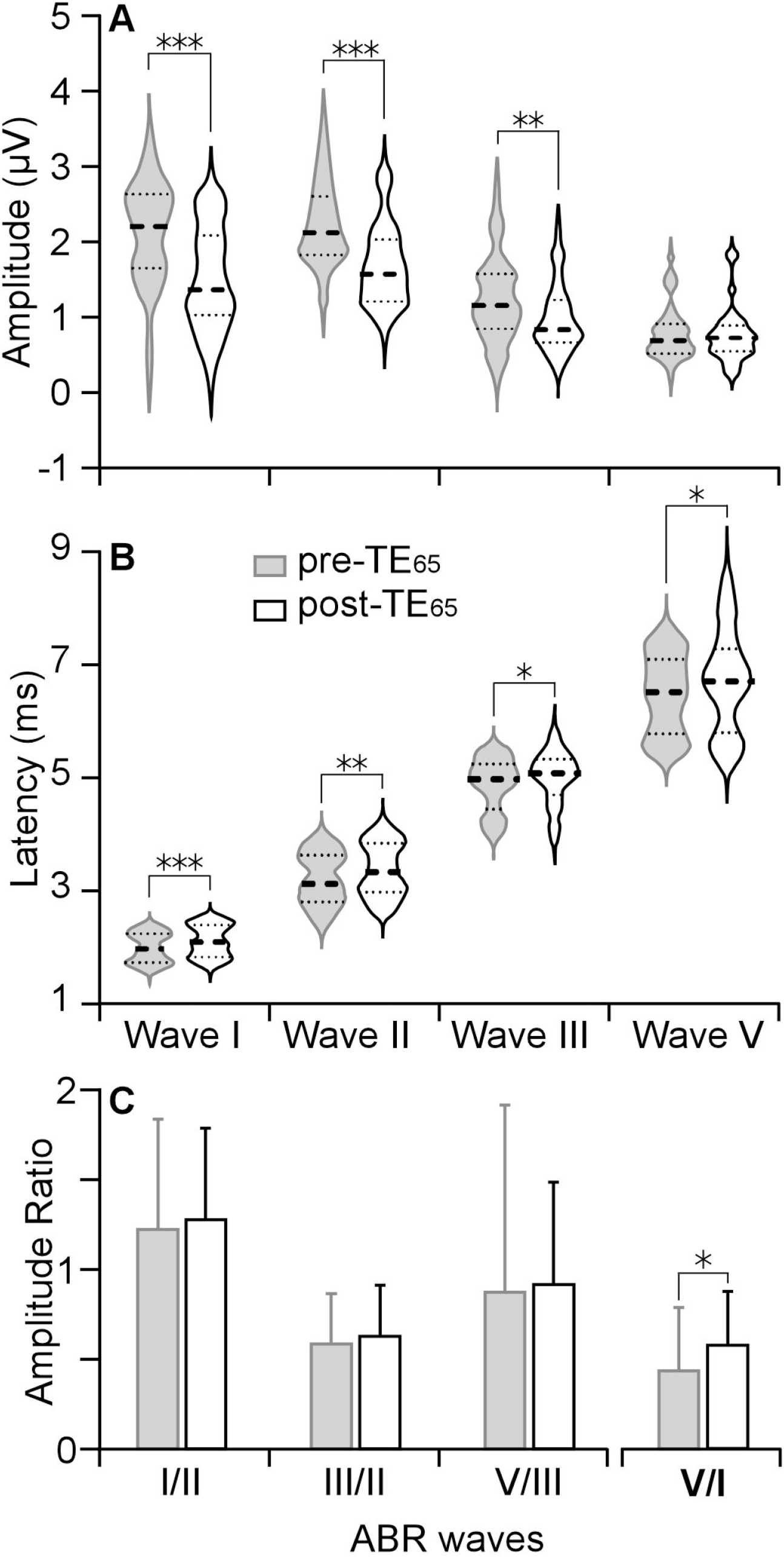
Comparison of the TE_65_-induced changes in the amplitudes and latencies of the Waves I-V to the tone burst at the exposure frequency and 60 dB SPL. TE65 significantly decreased the amplitudes (**A**) and increased the latencies (**B**) of Waves I-III. A trend of amplitude increases from Wave I to Wave V was observed as well as a significant increase in the ratio of Wave V to Wave I (**C**).

The TE_65_-induced inter-wave changes were examined by the changes in the amplitude ratios between any two waves. The ratios demonstrated a trend of consistent increase from Wave I to Wave V and the ratio of Wave II to Wave I showed the largest increase. However, these increases were statistically insignificant (Fig. 4C). We therefore compared the amplitude ratios of Wave V to Wave I; this ratio represents a summation of the amplitude differences among all waves. As expected, the TE_65_ significantly increased the ratio of Wave V to Wave I (0.43 ± 0.35 vs. 0.57 ± 0.30, p < 0.01). The largest increase in the ratio of Wave II to Wave I may be due to the significant decrease in the absolute amplitude of Wave I while the increases among the other waves might signify the compensatory enhancement of the activities in the neural structures following the response of the auditory nerve.

### Degraded correlations of amplitude and latency changes from Wave I to Wave V

Although the response latency is always negatively associated with the response amplitude, we further examined the correlation of the TE_65_-induced changes in amplitude and latency of all ABR waves. In Wave I, a decrease in amplitude was always accompanied with an increase in latency. The change in latency was negatively and linearly correlated to the change in amplitude (Fig. 5A, p < 0.001). This high correlation was maintained in Wave II except for two examples (Fig. 5B, p < 0.001). More exceptions were also observed in Wave III and Wave V; our increased samplings did not follow the rule of the decreased amplitude accompanying with increased latency or vice versa (Figs. 5C and 5D). Because of increased exceptions, the TE_65_-induced changes in the amplitude and latency became uncorrelated in Wave III and Wave V (p > 0.05). The insets in Figure 5 use vectors to better visualize the degraded correlation of the changes in amplitude and latency from Waves I through V.

**Figure 5.**
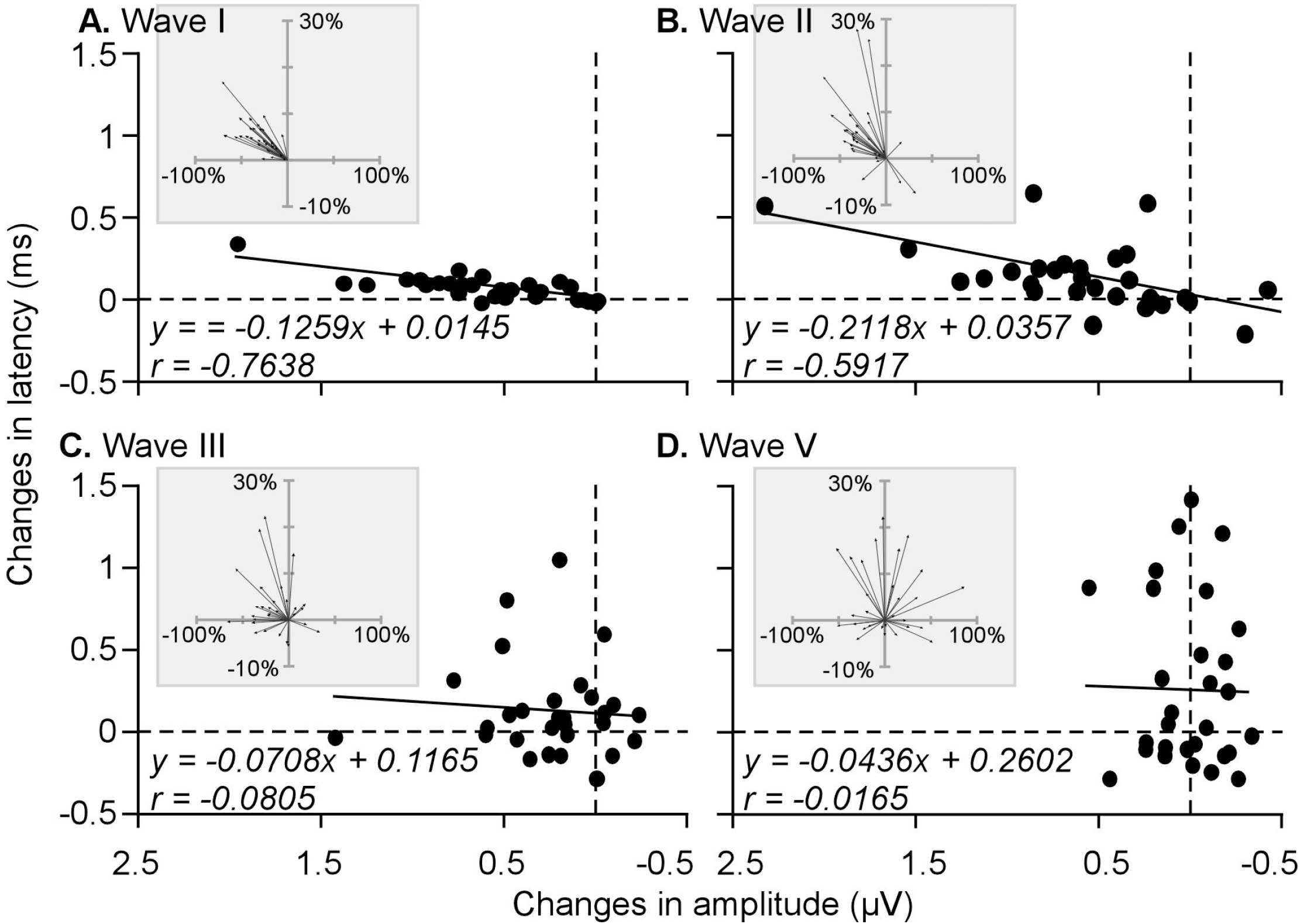
Correlations of TE_65_-induced decreases in the amplitude and increase in the latency of ABR waves. High correlations were observed in Waves I (**A**) and II (**B**) and poor correlation were observed in Waves III (**C**) and V (**D**). **Insets:** vectors show the percentage changes in the amplitude and latency.

### Long lasting effect of TE_**65**_

The effects of the TE_65_ on the ABR were not transient but last for hours. To quantify the recovery course of TE_65_ effects, the ABR threshold and Wave I amplitude were recorded in 10 mice for 6 hours once tone exposure was stopped. As shown in Figure 6, the maximal effects were immediately reached after TE_65_, including the increase in ABR threshold and decrease in Wave I amplitude to a tone at 60 dB SPL. On average, the threshold was increased by 6.11 ± 2.20 dB (p = 0.0002, n = 10), and the wave amplitude was decreased by 27.20 ± 10.80% (n = 10, p < 0.0001). The changes were equivalent to those calculated on 30 samples as presented above. The extent of TE_65_- induced changes gradually declined over time. The threshold levels dropped at a rate of approximately 1.019 dB/hr and were close to the pre-TE_65_ levels (19.44 ± 5.27 dB SPL vs. 16.67 ± 3.54 dB SPL, p = 0.054) at 6 hours after tone exposure. Since the threshold levels were significantly higher than pre-TE_65_ levels for 3 hours after tone exposure, we can conclude that the threshold increases persisted for this length of time. The TE_65_-induced decrease in Wave I amplitude at 60 dB SPL underwent a similar course of 3 hours after tone exposure and the amplitude decreases were statistically significant during this period. At 4 hours after tone exposure, Wave I amplitude returned to and even slightly exceeded the pre-TE_65_ amplitude. In conclusion, the effects of TE_65_ on threshold and Wave I amplitude shared a similar recovery time course of three hours although the threshold recovery appeared even more delayed.

**Figure 6.**
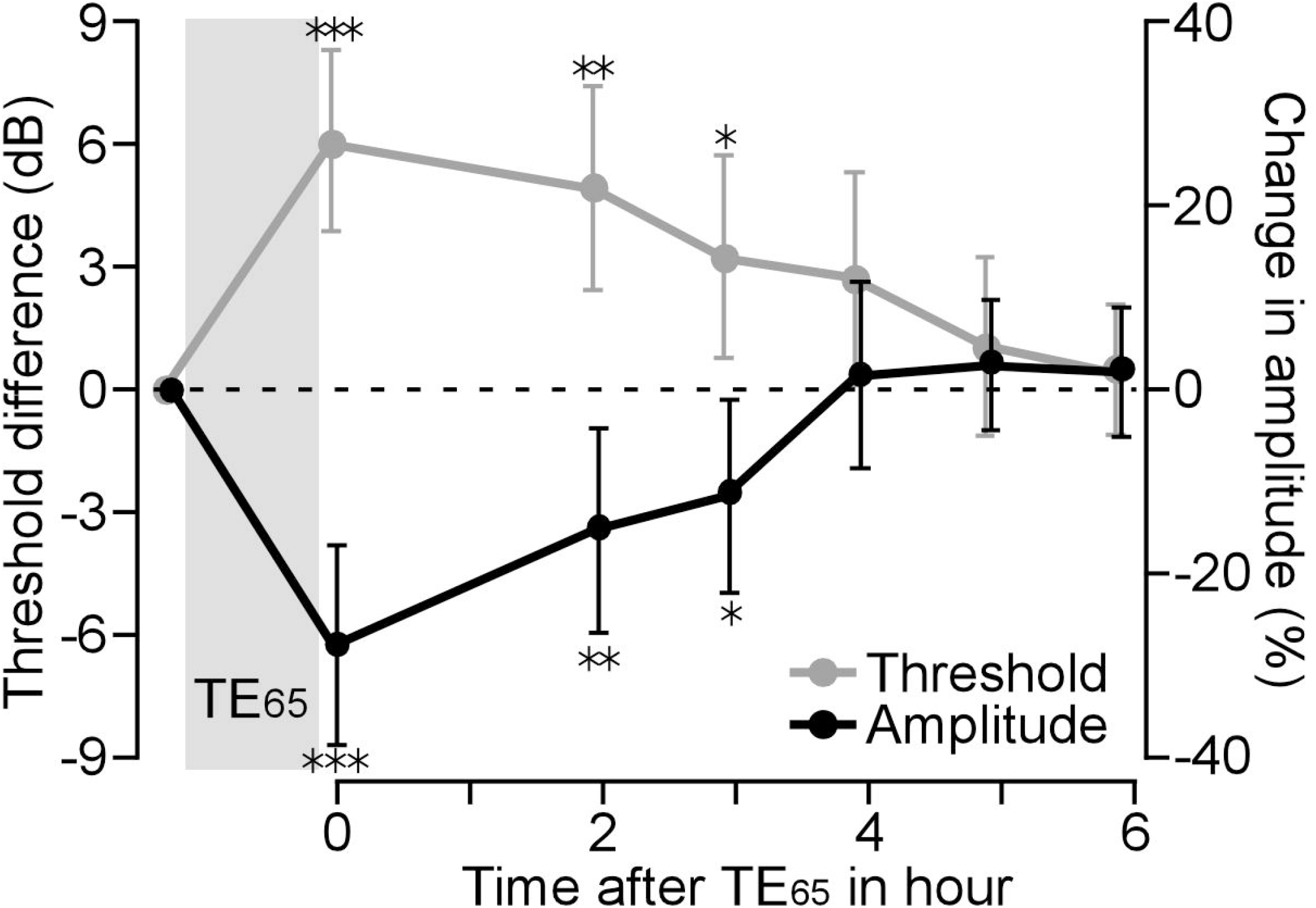
The time course of the changes in ABR threshold and Wave I amplitude after TE_65_. For both ABR threshold and Wave I amplitude, TE_65_-induced changes lasted at least 3 hours after the exposure; these changes remained statistically significant.

## Discussion

Our data show that TE_65_ increased ABR threshold by up to 15 dB according to the measurements of the greatest ABR waves (typically Waves I/II). Importantly, the threshold increase was specific to the frequency of the exposed tone (Fig. 2). Along with the threshold increase, a significant decrease in amplitude and increase in latency of ABR Waves I-III were also observed; the TE_65_-induced change in response to the tones was greatest in the 45-60 dB SPL range (Fig. 3C) and lasted for at least 3 hours. Notably, the amplitude decrease was negatively correlated with the latency increase in Wave I. This correlation gradually degraded from Wave I through to Wave V (Fig. 5). The effects of the TE_65_ on the ABR threshold and Wave amplitude lasted for at least 3 hours (Fig. 6).

### Input-specificity of TE_65_-caused increase in ABR threshold

Several studies using moderate narrow-band noise demonstrate that the auditory responses of cortical and subcortical neurons are altered or impaired mostly within the frequency range of exposure noise (Pienkowski and Eggermont, 2012; Thomas et al., 2019). To characterize the frequency-specific effect, this study employed pure tone as the acoustic exposure so that the exposure energy can focus on a single frequency. Our ABR data show that the significant threshold elevation was in a range of exposure frequency to the frequencies 0.5 octaves above while the change in thresholds to the frequencies below the exposure frequency was minimal. This suggests that the impact of tone exposure is restricted at the exposure frequency and its neighbouring high frequencies. As is well-known, sound-triggered vibration of the basilar membrane travels from the base (high frequency) toward the apex (low frequency). The displacement of a vibration gradually increases until it reaches a maximum at the segment corresponding to the sound frequency and sharply dissipates afterwards (Sohmer, 1997; Pickles, 2015). This theory of a sound wave travelling along the basilar membrane in the cochlea may well describe the frequency-specificity of TE_65_-caused increases in ABR threshold. Due to the inherent limitations of ABR, further clarification is required by directly measuring inner ear function such as compound action potential (CAP) following TE_65_.

### Possible mechanisms underlying ABR threshold increase

Since ABR Wave I primarily represents the tone-evoked neural activity of the auditory nerve terminals in the inner ear, our analysis of the decreased amplitude and prolonged latency of Wave I indicate that TE_65_-caused ABR threshold increases mainly originate from impairment in the sensory transduction in the cochlea. This consequence is further supported by the evidence that the exposure of rats to a narrow-band noise of 65 dB SPL significantly decreases CAP amplitude and increases CAP thresholds (Liu et al., 2021).

It is known that hearing loss caused by high and moderate noise exposure can result in the dysfunction of outer hair cells, inner hair cells or both in the cochlea (Le Prell, 2019). Outer hair cells, serving as a biological amplifier, play a crucial role in hearing sensitivity (Ashmore, 2008; Santos-Sacchi and Navaratnam, 2022) and their function can be indirectly measured by otoacoustic emission. Inner hair cells, serving as a transducer, convert mechanical energy of basilar membrane vibration to electrical energy of depolarized membrane potential, which evokes action potentials of the auditory nerve via the synapses (ribbon synapses) of inner hair cells on peripheral branches of spiral ganglion cells (Wichmann and Moser, 2015; Wagner and Shin, 2019). It is generally believed that the exposure to loud noise can destroy hair cells and that even moderate noise (e.g., 100 dB SPL) leads to the impairment of hair cells including the loss of ribbon synapses (Kujawa and Liberman, 2009; Lin et al., 2011). These impairments are directly associated with the decline in hearing (Liu et al., 2019; Fernandez et al., 2020). Gannouni and colleagues (2015) provided a more detailed understanding of inner ear damage in rats that were exposed to narrow-band noises of either 70- or 85-dB SPL (6 hr/day for 3 months). Under transmission electron microscopy, obvious deformation/destruction of the Organ of Cortis is evidenced. These noise-induced damages include but are not limited to stereocilia fusion/disorganization, cell body perforation, loss of cell organelles and cytoplasmic vacuoles in both outer and inner hair cells in limited basilar membrane segments that are likely due to frequencies carried in the exposed noise. The number of spiral ganglion cells is also decreased following noise exposure. It is worth noting that these impairments are more serious in rats exposed to 85 dB SPL as opposed to 70 dB SPL. The functional impairment of outer hair cells is also evidenced by the reduction of the distortion product otoacoustic emission in rats that are exposed to narrow-band noise at a level of 68 dB SPL for a week (Zhao et al., 2018).

Taken together with these facts, our study concluded that the exposure to a tone of 65 dB SPL for 1 hour may, to some extent, damage outer and inner hair cells as well as ribbon synapses but that the damage is milder and restricted in more limited basilar membrane segments corresponding to the exposed frequency and the neighbouring high frequencies.

### Functional alteration of the low brainstem

Due to impairments in the inner ear, altered inputs of auditory nerve to the brain can lead to functional changes in the central auditory system. Previous studies have documented that moderate noise (around 100 dB SPL) exposure largely changes the neural presentation and processing of auditory information in the auditory cortex (Wang et al., 2002; Pienkowski and Eggermont, 2009; Munguia et al., 2013; Auerbach et al., 2014; Eggermont, 2017). Although related knowledge in the inferior colliculus remains limited, it appears that prolonged exposure to moderate noise causes changes in auditory information processing from the cochlear nucleus to inferior colliculus (Brozoski et al., 2002; Hesse et al., 2016; Schrode et al., 2018; Palmer and Berger, 2019). However, exposure to low level noise (65 dB SPL for 4 weeks) does not cause detectable changes in the inferior colliculus even if a reduction of cochlear CAP amplitude is observed (Liu et al., 2021).

ABR studies allow assessment of the electrical events from the auditory nerve (Wave I) to the inferior colliculus (Wave V) simultaneously in one recording. Our data show a significant decrease in the amplitude of Waves I-III but not in the amplitude of Wave V. This is supported by previous investigations (Bakay et al., 2018; Schrode et al., 2018). The novel finding in this study, by analyzing the change in amplitude and latency of different waves, is the degraded correlation of the amplitude vs. latency change from Wave I through to Wave V (Fig. 5). This degraded correlation suggests that quick neural compensation or adaptation may occur in the low brainstem in response to the TE_65_-induced peripheral damage. Considering that ABR sampling was performed immediately after 1-hour tone exposure in this study, it would be interesting to investigate in what form and how the degraded correlation of amplitude vs. latency of Wave I through to Wave V is presented when tone exposure is repeated or extended for days, weeks and months.

### Clinical implication for HHL diagnosis

Normal hearing has a range from 0 dB HL to 20 dB HL and, according to WHO standards, hearing loss is defined as a diagnostic hearing threshold of higher than 20 dBA (Chadha et al., 2021). HHL is diagnosed because an individual presents with hearing difficulties but has a normal hearing threshold on routine audiograms. One possibility here is that individuals with HHL have an increased hearing threshold that still falls within the normal range based on current clinical guidelines.

Different from the over 20 dB threshold shift in response to loud noise exposure (Rüttiger et al., 2013; Jensen et al., 2015; Amanipour et al., 2018; Skoe and Tufts, 2018), the moderate-noise-caused threshold increase is relatively subtle, e.g., 7 dB in our study or 8 dB in Liu’s study (Liu et al., 2021). Therefore, it is not surprising that audiologists inform individuals with complaints of mild hearing impairment (e.g., a few dB increase) that their hearing is ‘normal.’ Our findings shed a different light on a possible approach to mild hearing loss diagnosis in HHL patients. A solution may be the regular use of ABR studies i.e. yearly, in individuals with hearing difficulties with prolonged exposure to even a mildly noisy environment. Audiologists would gain a more accurate diagnosis of patient hearing by comparing the ABR thresholds over time as well as the degraded correlation of amplitude vs. latency from Wave I to Wave V.

## Summary

Our study demonstrates that the exposure to a pure tone for 1 hour, which corresponds to the low levels of a normal conversation, may led to significant changes in the ABR in a frequency-specific manner. These ABR findings would help to confirm some degree of impairment in the inner ear. From this, we can infer that ‘safe’ sound may not be necessarily safe but can potentially increase hearing threshold and change auditory information processing in the brain. Furthermore, our findings provide valuable information for the consideration of new approaches for audiologists to exam the hearing of individuals who experience hearing difficulties despite having hearing thresholds that fall within the range of normal hearing.

## Acknowledgment

This work is supported by the Natural Sciences and Engineering Research Council of Canada (DG261338-2009) and the Rozsa Endowment for Auditory Research award from the Cumming School of Medicine, University of Calgary.

